# Myxomycetes have an allorecognition system that can only discriminate between intraspecies

**DOI:** 10.1101/2024.02.15.580440

**Authors:** Mana Masui, Phillip Yamamoto, Nobuaki Kono

**Affiliations:** Faculty of Environment and Information Studies, Keio University, Fujisawa, Kanagawa, 252-0882, Japan; Graduate School of Media and Governance, Keio University, Fujisawa, Kanagawa 252-0882, Japan; Institute for Advanced Biosciences, Keio University, Tsuruoka, Yamagata, 997-0017, Japan

**Author notes:** **Author for correspondence:** Nobuaki Kono.

**Keywords:** Slime mold, Myxomycetes, Plasmodium, Allorecognition

## Abstract

Myxomycetes are multinucleate unicellular organisms. They form a plasmodium that moves by protoplasmic flow and prey on microorganisms. When encountering intraspecifics, the plasmodium has the capacity for ‘fusion’, actively approaching and fusing its cells, or ‘avoidance’, altering its direction to avoid the other individual. This is an allorecognition ability. However, it remains unclear whether the range of allorecognition extends to other species, and its ecological significance is also obscure. Here, we conduct a quantitative evaluation of contact responses from closely related species of plasmodium to clarify the recognition range of the allorecognition system in Myxomycetes. Behavioral assays demonstrate that the allorecognition system recognizes individuals within the same species while failing to recognize those of different species. The allorecognition is an extremely narrow and inward-focused mechanism, arguing for a highly specialized system of self-other recognition.

**Summary statement:** Myxomycetes plasmodium can recognize each other only if the other individuals they encounter are of the same species.

## INTRODUCTION

Myxomycetes is multinucleate unicellular organisms belonging to the supergroup Amoebozoa [1,2,3]. They possess up to 800,000 nuclei per 1 mm^3^ [4]. Approximately 1,200 species of these Myxomycetes have been described worldwide [5], exhibiting various morphological forms throughout their life cycle, including the trophic form, fruiting body, and slime mold amoeba [4, 6]. Nutrient uptake in plasmodium is accompanied by migration under the action of protoplasmic flow at a rate of 1 mm per second, a process driven by the cytoskeletal elements of actin and myosin [7]. The primary nutritional sources, comprising chiefly of bacteria and fungi, are assimilated via endocytosis [8].

The cellular growth strategy during the plasmodial phase is notably intriguing. Beyond growth via synchronous nuclear division occurring in consistent cycles [4], plasmodia also employ a fusion strategy with other individuals of the same species [9, 10, 11]. This phenomenon has been reported in the genera *Physarum* and *Didymium* [12, 13].

Fusion amongst plasmodia is not mediated by nuclear fusion as in sexual reproduction, but is a phenomenon of sharing cytoplasm while retaining individual nuclei [9]. Fusion is not universally feasible across all plasmodial relations. Upon interspecific encounters, plasmodia recognize intraspecifics through ‘allorecognition’ and facilitate a choice between ‘fusion’, in which they actively approach and fuse, and ‘avoidance’, in which they change direction to avoid the other individual (Fig. 1A,B) [14].

**Fig. 1.**
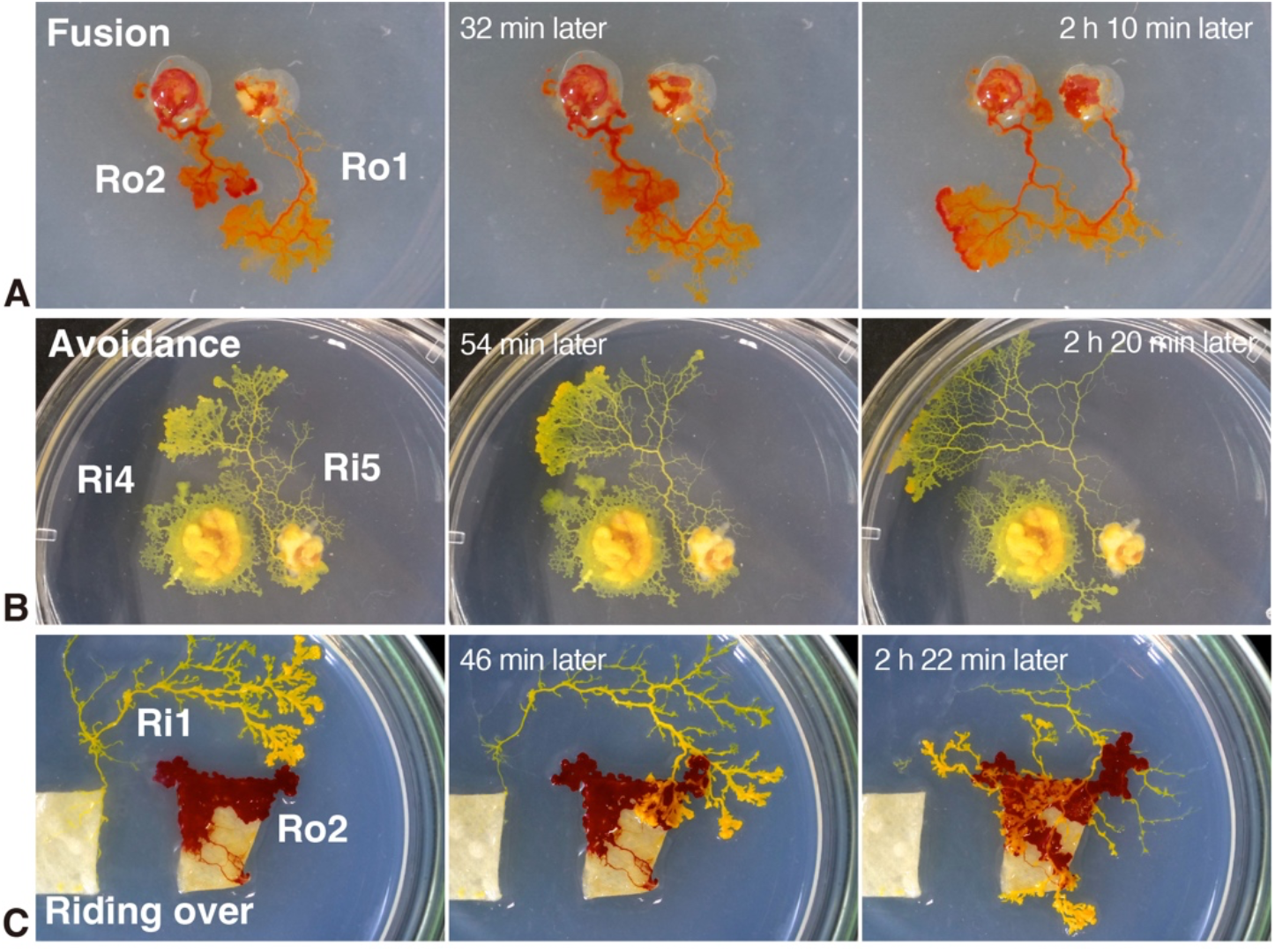
These pictures demonstrate typical behavioral examples. (A) Fusion behavior: after two strains of *P. roseum* (Ro2 and Ro1) encounter, they fused with each other and became one individual. (B) Avoidance behavior: after two strains of *P. rigidum* (Ri4 and Ri5) came into contact with each other, Ri5 changed the direction of movement and avoided without fusion. (C) Riding over behavior: yellow *P. rigidum* (strain Ri1) riding over the red *P. roseum* (strain Ro2) without showing any reaction.

The genetic loci implicated in this selective process have been hypothesised, although their precise roles remain obscure [15, 16, 17]. The ecological relevance of such allorecognition behavior remains to be fully understood. In related studies, a cellular slime mold, which shares a close evolutionary relationship with Myxomycetes. The cellular slime mold is known to gather amoebae of the same species to form reproductive structures, such as fruiting bodies and spores, which use chemical signals to attract other species [18]. The formation of the fruiting bodies takes place only in amoeboid cells of the same genetic lineage [19]. This suggests that the allorecognition behavior in cellular slime molds help them to find intraspecifics, a critical factor for the survival of their offspring.

Ants, as eusocial insects, process the ability to recognize their nest mates through cuticular hydrocarbons. This mechanism can also identify different species and non-nest mates of the same species [20, 21, 22]. This recognition behavior in ants contributes to the survival of the colony as a whole by discriminating between the elimination of external enemies and the communication of information with nest mates.

Considering the examples described thus far, the following hypothesis can be posited regarding the ecological role of the allorecognition. Should the allorecognition system encompass Myxomycetes broadly, extending to interspecies recognition, it might function as a strategy for mitigating competition, for instance, in nutritional aspects, with individuals beyond the fusion capability. Conversely, if the system is confined to intraspecies recognition, it would be insufficient in circumventing competition from other plasmodia. Hence, this system could be construed as optimally designed for searching fusible individuals.

Here, we aim to determine the adaptive range and the ecological significance of the allorecognition system by examining how Myxomycetes plasmodium responds to closely related species.

Behavioral test utilize *Physarum ridium*, a Myxomycetes already established in recognizing intraspecific plasmodium [12, 14]. While *P. ridium* is documented to engage in fusion and avoidance behaviors within same species, its reactions towards different species remain unexplored. In this context, *Pysarum roseum*, a different species within the same genus as *P. rigidum*, is introduced. By encountering *P. rigidum* with *P. roseum*, we investigate how far the allorecognition adapts, exploring its range of response.

It is also possible that an exclusive behavior that kills each other when plasmodia come into contact may be observed. This was also confirmed through behavioral test.

## RESULT

### Intraspecies behavior test

All of the anticipated encounter responses (fusion, avoidance, and no response) were observed in the Intraspecies behavior test, but their proportions varied widely (Fig. 1). Using *P. rigidum*, as the intraspecies behavior test, a total of 173 encounters. The results showed that 12 (6.94% of the total) exhibited fusion behavior. Avoidance behavior was observed in 154 cases (89.02% of the total). On the other hand, riding over behavior (no response) was observed in only 7 cases (4.05% of the total) (Fig. 2). Using *P. roseum*, a total of 39 encounters were observed. Unlike *P. rigidum*, no behavior of riding over on the partner was observed (Fig. 2). 17 (43.59%) of the total number of encounters showed fusion behavior, and 22 (56.41%) of the total number of encounters showed avoidance behavior (Fig. 2).

**Fig. 2.**
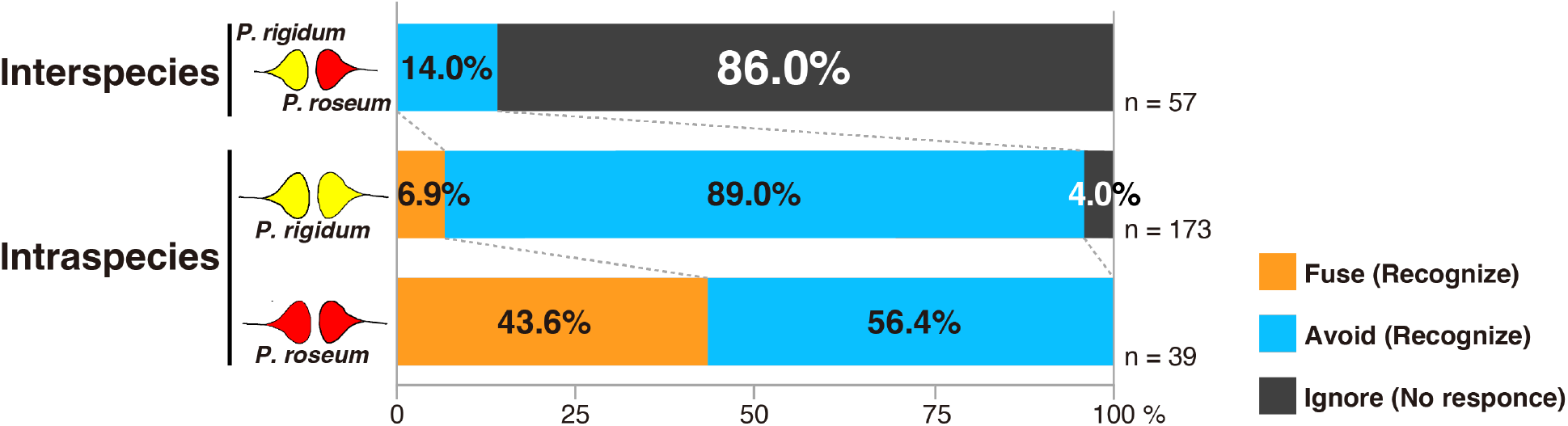
Assessment results for behavior tests between interspecies and intraspecies (*P. rigidum, P. roseum*). Ignore means the riding over behavior. There is a predominant difference in the evaluation results for each combination, with a tendency to ignore instead of the known allorecognition behavior between interspecies (χ2 = 231.357, p < .0001, Pearson’s chi-square test).

### Interspecies Behavior Test

The interspecies recognition test was conducted by confronting *P. rigidum* and *P. roseum*. A total of 57 encounter occasions were observed. Interestingly, quite unlike intraspecies behavior test, fusion behavior was never observed. The avoidance behavior was exhibited in eight cases or 14.04% of the total. The remaining 49 cases, or 85.96% of the total number of cases, exhibited riding over behavior (Figs. 1C and 2).

### Comparison of the percentage of allorecognition behaviors

The results of the behavior test evaluation (fusion, avoidance, and no response) of all 269 encounters were compared between different species/same species *(P. rigidum* and *P. roseum*), and significant differences were found (χ2 = 231.357, p < .0001, Pearson’s chi-square test) (Fig. 2). This result indicated that the slime molds tend to not show fusion and avoidance, known as allorecognition behavior, with respect to other species. In other words, they did not specifically recognize other species and ride over them as obstacles. In all encounters, the plasmodium survived for more than 24 hours after Myxomycetes decided their behavior (fusion, avoidance, and no response).

## Discussion

### Discovering a third behavior during plasmodium encounters

This study is the first report that *P. roseum* has the ability to recognize self and others and to fuse, confirming that self and others recognition behavior is a widely common behavior in the genus *Physarum*.

Furthermore, fusion and avoidance, which are allorecognition behaviors, were rarely observed between interspecies (Fig. 2), even though they belong to the same genus *Physarum*.

Cross-genus verification of interspecies recognition has previously been done, with the *Physarum* and *Fuligo*, no cytoplasmic uptake on contact has been observed. [27].

The fact that no interspecies fusion behavior at all is observed in our experiments confirms previous studies.

In addition to this, a behavior that could be interpreted as completely ignoring the other individual was prominently observed in interspecies. It is a behavior that neither fuses nor avoids but rides over other individuals (Fig. 2).

When the opponent is a homologous heterologous individual, if it is a target that cannot fuse, the behavior of avoiding the opponent has been observed in most cases [14].

Avoidance behavior is a behavior in which the target is once recognized as a Plasmodium of a different individual and judged that can’t become a single individual by fusion.

However, the behavior of ignoring is considered to be a failure to recognize the target as a Plasmodium of myxomycetes in the first place. Therefore, this behavior can be defined as a third response that a plasmodium takes when it comes into contact with other individuals.

As well as no fusion, no behavior has been observed in previous studies and this study in which heterophiles attack each other on contact.

Examples of ignoring between interspecies without attacking each other can also be seen in the field (Fig. 3). This means that Myxomycetes have the opportunity to come into contact with many different species of Plasmodium. From the above, it can be said that contact with different species of plasmodium is not completely unexpected for plasmodium and that they are not recognized as objects of fusion or avoidance and do not actually harm each other.

**Fig. 3.**
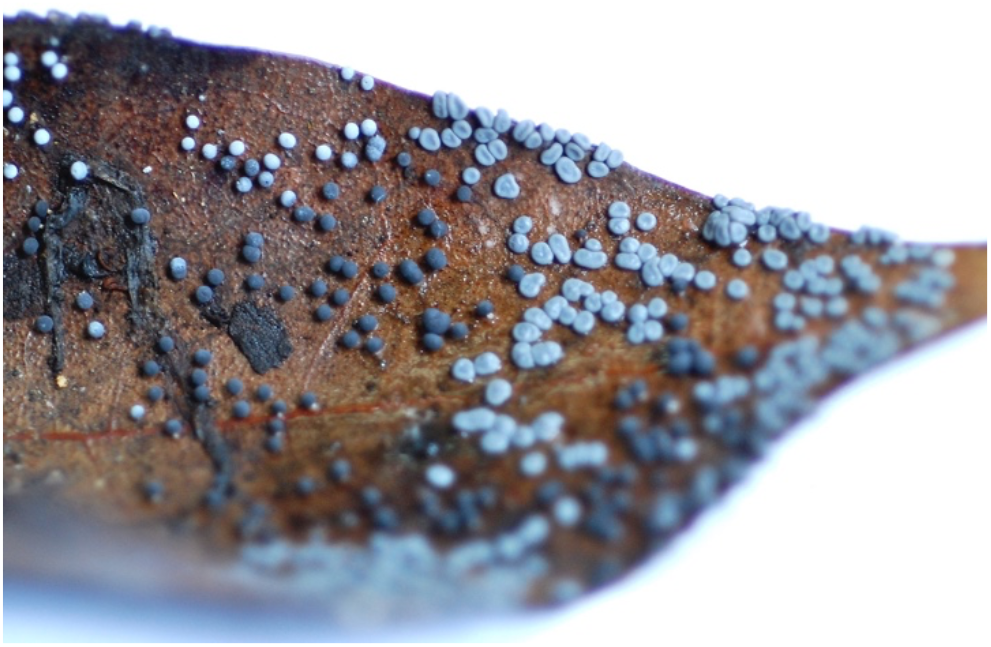
Fruiting bodies are formed by two species of Myxomycetes next to each other. The grey, spherical one is *Diderma effusum*. The dark grey, irregularly elliptical one is *Didymium squamulosum*.

As they both formed complete fruiting bodies, it can be said that they did not fuse or attack each other in the plasmodium stage.

### The plasmodium’s allorecognition system can discriminate only between intraspecies

The behavior of ignoring found in this experiment was almost never observed in encounters within the intraspecies and was therefore interspecies-specific. This indicates that the allorecognition system of plasmodium can only recognize plasmodium of the same species. Based on these results, it can be concluded that this system is insufficient for avoiding plasmodium of other species and that it is suitable for searching for plasmodium of the same species, a group with the potential to fuse with each other. However, if the significance of this system is considered as above, it is necessary to consider its significance in the plasmodium of behavior ‘fusion’. In spite of this, to date, the benefits of fusion have not been clearly demonstrated and have hardly been discussed in previous studies. Fusion between plasmodium is a phenomenon in which the membranes of the cells dissolve and the protoplasm mixes with each other but does not fuse with the nuclei. It is therefore unlikely to contribute to increased genetic diversity in reproduction. One change that can be stated with certainty for plasmodium in fusion is the increase in the size of their own cells as a result of incorporating different individuals. The advantage of this is the formation of more fruiting bodies. Myxomycetes form dozens to hundreds of fruiting bodies from a single plasmodium, in which tens of thousands to millions of spores are present [28]. The number of spores produced per fruiting body varies among species to some extent, but the total number of spores formed by a single plasmodium is not species-specific. Instead, the total number of spores formed by a single plasmodium depends on its size. This implies that there is a clear advantage for the plasmodium in increasing its cell size, which increases the chances of producing more offspring. However, the benefits and drawbacks of plasmodium cell size enlargement during plasmodium including potential shifts in energy metabolism efficiency could not be examined due to inadequate knowledge.

However, significant variations in the response to contact between intra and interspecies could provide essential insights into inter-individual responses in Myxomycetes. For instance, previous research has indicated that the slime sheath produced by plasmodium is involved in allorecognition behavior. Signal substances for recognition are yet to be identified. The main component of slime sheath has been identified as a glycoprotein, but the other components continue to be debated. This study suggests that the signal substance used here is species-specific. This is because known allorecognition behaviors such as fusion and avoidance have only been observed within the intraspecies. The fact that neither of the two species used in the verification showed any interspecies response, despite the similarity of the allorecognition behavior within the species. It suggests that the mechanisms of allorecognition are similar, but the signaling molecules are different.

Although this study has allowed us to consider the ecological significance of the allorecognition behavior of Myxomycetes, still unknown about the molecular mechanisms. We believe that clarifying the details of the loci that determine whether fusion is possible or not and the mechanisms that control the behavior will bring us closer to understanding the full range of interactions between individuals in Myxomycetes.

## MATERIALS AND METHODS

### Samples

Sampling was carried out in Saitama Prefecture, Japan, using seven independent *P. rigidum* (Ri1∼7) and three *P. roseum* strains (Ro1∼3). Species identification was performed by morphological characteristics of their fruiting bodies [23, 24] or by DNA barcoding. PCR primers for the SSU rRNA region of DNA barcoding were phf1b (5’-AAAACTCACCAGGTCCAGAT-3’) for forward and phr2b (5’-TACAAAGGGCAGGGGACGCAT-3’) for reverse. PCR conditions following previous studies 94°C, 1 min; [98°C, 10 sec; 55°C, 15 sec; 68°C, 1 min] x 30; 68°C, 5 min [25]. Sanger sequencing was performed by DNA Sequencing Service (Eurofine Genomics Ltd.). The sequence data were organized using MAFFT alignment [26] and then utilized as a query in species identification through NCBI BLAST.

Plasmodium were cultured on 2% agar in petri dish of 9 cm in diameter and fed with water and oat flakes as a nutrient source every day. Temperature conditions were maintained at 25°C for *P. rigidum* and 22°C for *P. roseum* using a cool incubator, while shading.

### Behavior test

Behavior tests to verify the allorecognition behavior of plasmodium were conducted based on a previously established method [14]. Two plasmodia were placed 3 mm apart on a 2 % agar medium and recorded by time-lapse photography every minute until the two individuals fused or two days had elapsed. The experimental environment was kept at 25°C, with no direct light exposure, and watering with sterile water to prevent the surface of the medium from drying out. Both *P. rigidum* and *P. roseum* were used in intraspecies and interspecies behavior tests.

Behavior tests were qualitatively assessed using a tripartite framework comprising ‘fusion’, ‘avoidance’, and ‘no reaction (ignore)’. These indicators were meticulously documented and categorized up to the point of either ultimate fusion or total cellular separation. Encounter cases were classified using time-lapse photography. Fusion was defined as the fusion of cells with each other. Avoidance is defined as a sop in movement or a change in direction of travel without fusion occurring during observation. If the cells did not exhibit either of these two behaviors and also exhibited the behavior of riding over on the other individual, it was classified as ‘no response (ignore)’.

The above three indices were cross-tabulated among different species/species (*P. rigidum, P. roseum*) for all encounters in the behavior test. Pearson’s Chi-squared test was performed on them. JMP (version 14.3) was used for statistical analysis.

## Acknowledgement

We thank the Saitama Midori-no-mori Nature Park for their cooperation in sampling the strains and Maaya Domon for providing technical support.

## Footnotes

### Author contributions

Conceptualization: M.M., N.K.; Methodology: M.M., N.K.; Validation: M.M., P.Y., N.K.; Formal analysis: M.M.; Investigation: M.M.; Resources: M.M., N.K.; Data curation: M.M.; Writing - original draft: M.M.; Writing - review & editing: M.M., P.Y., N.K.; Visualization: M.M.; Supervision: P.Y., N.K.; Project administration: N.K.; Funding acquisition: M.M., N.K.

### Data availability

Raw data used in this research are available from the Dryad Digital Repository: https://doi.org/10.5061/dryad.cfxpnvxd9 [29].

